# Gestational and postnatal age associations for striatal tissue iron deposition in early infancy

**DOI:** 10.1101/2023.06.30.547249

**Authors:** Laura Cabral, Finn Calabro, Jerod Rasmussen, Will Foran, Luci A. Moore, Alice Graham, Thomas G O’Connor, Pathik D Wadhwa, Sonja Entringer, Damien Fair, Claudia Buss, Ashok Panigrahy, Beatriz Luna

## Abstract

Striatal development is crucial for later motor, cognitive, and reward behavior, but age-related change in striatal physiology during the neonatal period remains understudied. An MRI-based measure of tissue iron deposition, T2*, is a non-invasive way to probe striatal physiology neonatally, linked to dopaminergic processing and cognition in children and adults. Striatal subregions have distinct functions that may come online at different time periods in early life. To identify if there are critical periods before or after birth, we measured if striatal iron accrued with gestational age at birth [range=34.57-41.85 weeks] or postnatal age at scan [range=5-64 days], using MRI to probe the T2* signal in N=83 neonates in three striatal subregions. We found iron increased with postnatal age in the pallidum and putamen but not the caudate. No significant relationship between iron and gestational age was observed. Using a subset of infants scanned at preschool age (N=26), we show distributions of iron shift between timepoints. In infants, the pallidum had the least iron of the three regions but had the most by preschool age. Together, this provides evidence of distinct change for striatal subregions, a possible differentiation between motor and cognitive systems, identifying a mechanism that may impact future trajectories.

**Highlights:**

- Neonatal striatal tissue iron can be measured using the T2* signal from rsfMRI
- nT2* changed with postnatal age in the pallidum and putamen but not in the caudate
- nT2* did not change with gestational age in any of the three regions
- Patterns of iron deposition (nT2*) among regions shift from infancy to preschool

## 1.0 Introduction

Rapid brain development takes place from the prenatal period into the first months of postnatal life, with contributing genetic and environmental influences that set up a template to impact future developmental trajectories. Critical for later cognitive, motor and reward-based behavior is the maturation of the striatum, which receives the primary glutamatergic and dopaminergic inputs for the basal ganglia, influencing behavioral responses^1^. Thus, investigating the early development of its neurophysiology can elucidate critical components that can influence later development. A non-invasive approach to probe changes in striatal neurophysiology in early development is through measuring tissue iron deposition^2–5^.

Brain tissue iron is stored as ferritin by mitochondria and accrues primarily in the basal ganglia^6, 7^. Ferritin is necessary for cellular respiration^8^ and energy production^9, 10^,. Importantly, tissue iron is located in the dendrites of dopamine neurons and has been found to co-localize with dopamine vesicles^11^ contributing to monoamine synthesis, including dopamine production^12, 13^.

MRI measures of striatal tissue iron have been associated with dopamine availability, and various MRI sequences have demonstrated their potential to measure striatal tissue iron developmentally^14, 15^, characterizing individual trajectories related to both cognitive and reward performance in adolescence^16–18^.

Although there is evidence that MRI-based striatal iron has the ability to identify developmental change in striatal neurophysiology, little work has extended this approach to neonates. Work linking iron to behavior initially focused on serum iron. Maternal iron availability can diminish iron stores in the fetus, where even mild iron deficiency has negative consequences^19, 20^. For example, infants who had low cord ferritin at birth were found to have lower scores on motor and cognitive tests at 5 years of age^21^. Infants who experience postnatal anemia also often have lower scores on similar motor and cognitive tests; these deficits have been seen in children up to 72 months^22–24^. Older children who were anemic in infancy have difficulty on tests of inhibitory control, set-shifting, and planning in later childhood^25, 26^.

Initial work linking iron availability to infant brain development showed children who were anemic in infancy had slower auditory brainstem responses and visually evoked potentials at 6-12 months and at 4 years^27, 28^. In older children from a similar cohort, decreased inhibitory control was related to longer N2 and P300 responses, which could reflect attention or abnormalities in frontostriatal connectivity^25^.

In anemic infants, deficits persist, despite iron therapy in infancy. Anemia causes a decrease in red blood cells and hypoxic conditions that may be visible on MRI^29^, and indicate behavioral deficits^30^. Hypoxia causes stored iron to be released from cells, leading to oxidative stress and iron-mediated cell death, increasing striatal iron content^31^. Hypoxia exerts diffuse effects, but striatal damage may be impactful because of the concentration and loss of dopamine neurons. Iron-mediated dopamine neurotoxicity is particularly harmful to surrounding cells^32^. Thus, altering early trajectories could lead to long-term deficits not reversible by simply resolving anemia.

Work in typically developing children has supported that striatal tissue iron is related to cognitive performance. In the caudate, often associated with cognition, striatal tissue iron was positively associated with working memory and spatial IQ in 6–11-year-olds^33, 34^. In 7-16-year-olds, basal ganglia iron deposition was related to greater processing speed and scores of general intelligence^35^. Despite evidence for meaningful relationships and the impact early life can have on striatal trajectories, little work has been done to characterize neonate or infant striatal iron deposition.

In foundational work^36^, Zhang et al. (2019) used two MRI-sequences, quantitative susceptibility mapping and R2*, to look at striatal iron from 12 neonates, with binned cross-sectional data into early toddlerhood. Across their sample, they found increases in overall striatal tissue iron between groups, demonstrating MRI based measures are sensitive enough to detect age-related change across the first two years of postnatal life. Ning et al (2014)’s work modeled change with infants with a postmenstrual age of 37-91 weeks, doing so separately for the caudate, putamen, and pallidum. They showed age-related associations across their sample, with increased tissue iron for all subregions into the first year of life. Ning (2019) extended this work to isolate early change, modeling when during the 6 years of postnatal life, linear relationships between age and iron emerged. With 28 infants under 6 postnatal months, the relationship between striatal iron and age was not evident until 3 months in the pallidum and putamen. In the caudate, this relationship was not apparent until 6 months^37^.

Previous work demonstrates striatal tissue iron can be measured in the first year and cross-sectionally changes with postnatal and postmenstrual age. Although these are interesting and established methods of characterizing age, solely accounting these only characterizes inter-individual variability with the age at scan, despite shifts in brain-trajectories when transitioning into the postnatal environment. Isolating the transition from prenatal to postnatal development remains unstudied. To assess this, it is necessary to model gestational age (GA) and postnatal age independently in linear models, allowing us to identify periods of increasing iron deposition that may be non-linear from conception to scan. If postnatal stimulation was driving iron deposition, we could see linear change with postnatal but not GA. Conversely, if access to maternal iron was driving deposition, linear change might be primarily associated with GA. If both related to iron similarly, we could expect to see linear relationships for both variables.

To assess the relationship between GA and postnatal age within striatal subregions, we use a dataset of 83 infants whose GA ranges from 34.57-41.85 weeks and postnatal age at the scan from 5-64 days. We quantify striatal tissue iron using a method previously used to characterize adolescent development, using the T2* signal from resting state fMRI scans. Tissue iron is paramagnetic and has a short transverse relaxation time (T2) in contrast with other tissues, making iron inversely related to T2*. We modeled T2* signal with gestational age and postnatal age independently to identify significant periods of age-related development in the three striatal subregions studied in previous work. In an exploratory analysis, we assessed how the distribution of striatal iron changes among striatal subregions longitudinally, in a subset of infants at preschool age.

## 2.0 Methods

### 2.1 Overall Sample

During pregnancy, maternal infant pairs were recruited to participate in a longitudinal study at the University of California, Irvine. Infants were excluded from the study if they met one of the following criteria: a congenital, genetic, or neurological disorder, were born before 34 weeks gestation, or there was maternal use of psychotropic medication or corticosteroids during pregnancy. Infants who met the inclusion criteria participated in an MRI scan neonatally. After successful preprocessing, we were left with 85 participants. One participant who contributed high motion data (Framewise Displacement >3) was removed from the analysis, leaving a total of 84 participants. An additional infant was removed because they did not have a recorded gestational age, leaving a total of 83 with complete data. Of the 83 who met criteria, infants had a mean gestational age of 39.25 weeks [Range 34.57-41.85] and a mean postnatal age of 24.94 days at the MRI scan [Range 5-64] at the MRI scan. When combining gestational age with postnatal age the infants had a mean postmenstrual of 42.81 weeks [Range 39.71-48.57].

Additional sample characteristics were gathered from the medical records, including obstetric risk, infant sex, and birth weight. Obstetric risk was a binarized outcome measure and was included as a two-factor variable in the models. Records indicating preeclampsia, hypertension, diabetes, severe anemia, severe infection, and vaginal bleeding indicated obstetric risk. Socioeconomic status was measured using parental self-report data, where maternal education was scored on a scale of 1-5. A subset of participants (N=26) took part in an additional MRI scan at preschool age (Mean Age: 5.31 years; Range: 3.37-7.06 years).

### 2.2 Infant MRI Scanning and Acquisition Parameters

As described in previous work utilizing this dataset ^38–40^, infants completed MRI scans after being fed while in natural sleep. A CIVO beaded head pillow was used, which wrapped around and under the body and head of the neonate, and the air was vacuumed out of the pillow, resulting in a firm swaddle that helped to immobilize the infant. Additional insulation from the pillow provided adequate hearing protection in combination with earplugs. Other descriptions of the infant scan protocol can be found in previous publications^41^.

Scans were then completed on a 3T Siemens Scanner (TIM Trio, Siemens Medical Systems Inc.) MPRAGE T1 scans (TR of 2400 ms, TE of 3.16ms, TI of 1200ms, voxels 1mm isotropic) with a Flip Angle of 8 degrees, a Matrix size of 256×256×160, with a total time of 6m18s were acquired. Additional T2 weighted scans were completed and used for segmentation (matrix size 256×256×160 with a TR of 3200 ms, and a TE of 255 ms, total time of 4m18s.) T2 scans had 1mm isotropic voxels and included a 0.5 mm slice gap.

Using gradient-echo planar imaging (EPI), a functional, resting state scan was acquired. These images were sensitive to the blood oxygen level dependent signal and were T2* weighted. The sequence had a TR of 2000 ms, a TE of 30 ms, and a field of view that was 220 x 220 x 160 mm, with a flip angle of 77 degrees. Full brain coverage was acquired by using a 1mm skip with 32 ascending-interleaved 4 mm axial slices. The first four volumes were discarded, after which magnetic field homogeneity was assumed. Infants completed resting state scans for a maximum of 195 volumes.

### 2.3 Overall Infant Anatomical MRI Processing

The processing for the structural data was obtained by running the <u>infant-abcd-hcp-pipeline</u> and extracting intermediary pipeline outputs in volume space. The infant-abcd-hcp-pipeline is derived from the anatomical minimal processing pipeline from the Human Connectome Project (HCP) (Glasser et al 2013). This HCP pipeline took the initial T1 and T2-weighted images and warped them to a desired template. The pipeline also created a binarized brain mask, corrected for artifacts and distortions, performed surface alignment, and registered data into a standard space.

The modified HCP pipeline used here was optimized for neonatal data, adding two initial steps. As neonatal and infant images have less space between the brain and skull and are often more susceptible to distortion than adults, ANTs DenoiseImage and N4BiasFieldCorrection were run to clean the images prior to HCP processing. After this, the pipeline followed the initial HCP workflow, running Pre-Freesurfer. Pre-Freesurfer provided the initial brain masks, segmentations, and created the warps between individual subject space and the chosen template. The procedure to create the first brain mask was optimized for infants, and both the T2 and T1 weighted images underwent a rigid body transform to align the ACPC with the infant MNI template, the infant template of choice in this study. The T2 image was then nonlinearly warped to the infant MNI template, and template brain mask was back transformed into individual subject T2 space. This mask was used to extract the T2 brain. Finally, a T1-weighted brain mask was derived by performing a rigid body registration between the T1 and T2 weighted images, and the transformed T2 brain mask was used to extract the T1 brain. The masked images were then non-linearly warped to the infant MNI template.

Segmentation can be difficult in neonatal images because of the reduced contrast between the grey and white matter. To ensure accuracy, additional steps were added to this stage in Pre-Freesurfer. 10 T1 atlases that had been manually segmented and labeled were non-linearly registered to tThe subject T1 images, already warped into in infant MNI space, were non-linearly registered to 10 T1 atlases that had been manually segmented and labeled. The ANTS Joint Fusion algorithm then calculated a correlation value between intensities for all voxels in each T1 and the atlases. Using the correlations, voxels were then defined as grey or white matter based on the majority consensus between the 10 atlases.

As in the HCP, the FreeSurfer pipeline was used next and served as a recon-all pipeline, with primary goals of segmenting the images into defined structures, reconstructing the white and pial cortical surfaces, and the registration of the processed images to a surface atlas, using FreeSurfer’s folding-based surface registration. Several modifications were made for the pipeline to be usableuseable in neonates. First, the adult FreeSurfer pipeline created surface images from the mean intensity of the preprocessed, native space images. Infant images have different intensities than adult images, so the intensity of the infant images was matched to the adults before undergoing this step. Finally, FreeSurfer’s image segmentation was replaced with a method developed by our OHSU including authors of this study. The segmentation used here defines white matter boundaries using a gradient descent algorithm, utilizing the adjusted intensity values in the infant images. After boundaries are identified, a white matter mesh is projected to the pial boundary and used to calculate midthickness. After this, FreeSurfer then used the normalized data to reconstruct surface images in native space.

PostFreeSurfer then took the restored images in native space and converted them into NIFTI and CIFTI images for use in the Connectome Workbench. These images were then transformed into atlas space. To optimize the pipeline for neonatal data, the transformation into atlas space was done using a non-linear registration with ANTS compressible fluid deformation, instead of the FNIRT registration that is used with the adults. This portion of the pipeline also produced the last brain mask and myelin maps.

### 2.4 BOLD Image Processing

Finally, the inputs from PreFreeSurfer, FreeSurfer, and PostFreeSurfer were used to run fMRIVolume to register striatal function and anatomy. The HCP fMRI Volume pipeline took the restored anatomical images and used them to register the functional data to the standard template. In this case, functional data was registered to the infant MNI atlas described above. The pipeline was modified to exclude distortion correction, since no field map was acquired in this sample. To complete the functional preprocessing, fMRIVolume carried out slice time correction, frame-frame realignment, and mode 1000 normalization. Framewise displacement (FD) was calculated at this stage. These preprocessing steps are consistent with previous work, and, importantly, do not contain global signal regression or bandpass filtering, as this would remove the global T2* signal that we are using to measure tissue iron.

### 2.5 Preschool MRI Acquisition Parameters

Children were scanned at the University of California Irvine with a Siemens 3T Prisma MRI scanner. As in adult studies, children were scanned while awake and wearing standard hearing protection. A T1 weighted MPRAGE scan (TR=2500 ms, TE=2.9 ms, TI=1070 ms, FOV=256 x 256 x 176 mm, with a flip angle of 8 degrees) was used to acquire the data. Voxels were 1 mm isotropic. As in the infants, a functional resting state scan (EPI), sensitive to the T2* signal, was acquired (TR=800 ms, TE=30 ms, FOV= 216 x 216 x 144, flip angle=52 degrees). Voxels were 2.4mm isotropic. Four children were scanned twice, brought back to gain additional data. When possible (N=2), we used the second scan for analysis. However, in N=2 cases, we used the first scan because, as identified with visual inspection, the second contained an artifact after nT2* processing.

### 2.6 Preschool Anatomical MRI Preprocessing

The overall preprocessing pipeline for the preschool children was modified from Hallquist et al. (2013). Bias correction was performed with FSL, and ROBEX was used to extract the brain from the image. Each participant’s brain was then warped using both a linear transformation (FLIRT) and non-linear transform (FNIRT) to the MNI ICBM152 09c template. It is easier to identify grey and white matter boundaries in children than in infants, and the child data both segmented and registered to the adult MNI template well. Brain masks were created based on the warps to the template, and from these FSL’s tissue type segmentation was used for boundary-based registration when aligning the functional images to the structural images.

### 2.7 Preschool BOLD Image Processing

As in Hallquist et al., (2013), functional images were motion and 4-D slice time corrected using FSL. They were skullstripped using FSL’s BET and were despiked using the Brain Wavelet toolbox in MATLAB. The functional images were then warped to the adult MNI template space using the warps derived from the processing of the structural images. As in the infant data, there was no distortion correction. These preprocessing steps are consistent with previous work in adolescents and adults using rsfMRI to estimate tissue iron, and, overall, as in the infant preprocessing described above. Most importantly, they do not contain preprocessing that would remove the overall T2* signal, such as bandpass filtering or global signal regression.

### 2.8 Calculation of Infant and Child Normalized T2*: A Measure of Striatal Tissue Iron

To measure tissue iron, the functional images, registered to template space were used to calculate normalized T2* (nT2*). Here, we were interested in quantifying a single measure of iron and not fluctuations that may occur in the rsfMRI signal across time. For each subject, nT2* was calculated on a per voxel basis, where each voxel in a volume was normalized to its volume’s whole brain median, making nT2* values relative to the rest of the brain. Having volumes that are normalized removes the variability in signal due to subject specific variables, such as inhomogeneity in the magnetic field due to subject placement and allows us to make between participant comparisons. Single values for each voxel were then computed by taking the median value for each voxel across the entire fMRI time course, resulting in a single, voxelwise nT2* image. Taking the median increases the stability of the measure.

Striatal regions of interest, the pallidum, putamen and caudate, were defined using atlases distributed with FSL. The pallidum was defined using the Harvard Oxford subcortical atlas (Desikan et al., 2006) and the putamen and caudate were defined using the Oxford-GSK-Imanova Structural and Connectivity Striatal atlas (Tziortzi et al., 2011). In adult space, there were two overlapping voxels between the putamen and pallidum. These were removed from the putamen ROI and were allocated to the pallidum, which was the smaller region. Regions were transformed into infant template space using ANTs non-linear registration. As nT2* was calculated in template space for all infants, the transformed Harvard-Oxford regions were used to localize the striatal regions, and mean values were extracted from each subject’s nT2* map. As iron has short transverse relaxation times, nT2* values are inversely related to tissue iron, where higher values indicate less tissue iron^17^.

## 3.0 Results

### 3.1 Overall Infant Regional Quantification

Gestational and postnatal age, as well as sex for the infants are described in the methods above and histograms that depict the distribution of gestational (**Supplementary Figure 1**) and postnatal (**Supplementary Figure 2**) age are found in the Supplementary Figures.

The caudate had the lowest mean nT2* signal (M= 1.014, SD= 0.033), followed by the putamen (M= 1.009, SD= 0.021) and the pallidum (M=1.029, SD=0.024). There were significant differences in nT2* between the regions, and paired sample t-tests showed the pallidum had significantly higher nT2* than both the caudate (*t*(82)=4.33, p<0.001) and the putamen (*t*(82)=16.75, p<0.001). There was no reliable difference in nT2* between the putamen and caudate (t(82)=1.51, p=0.136. The distribution for the three regions can be found in **Figure 1**.

**Figure 1.**
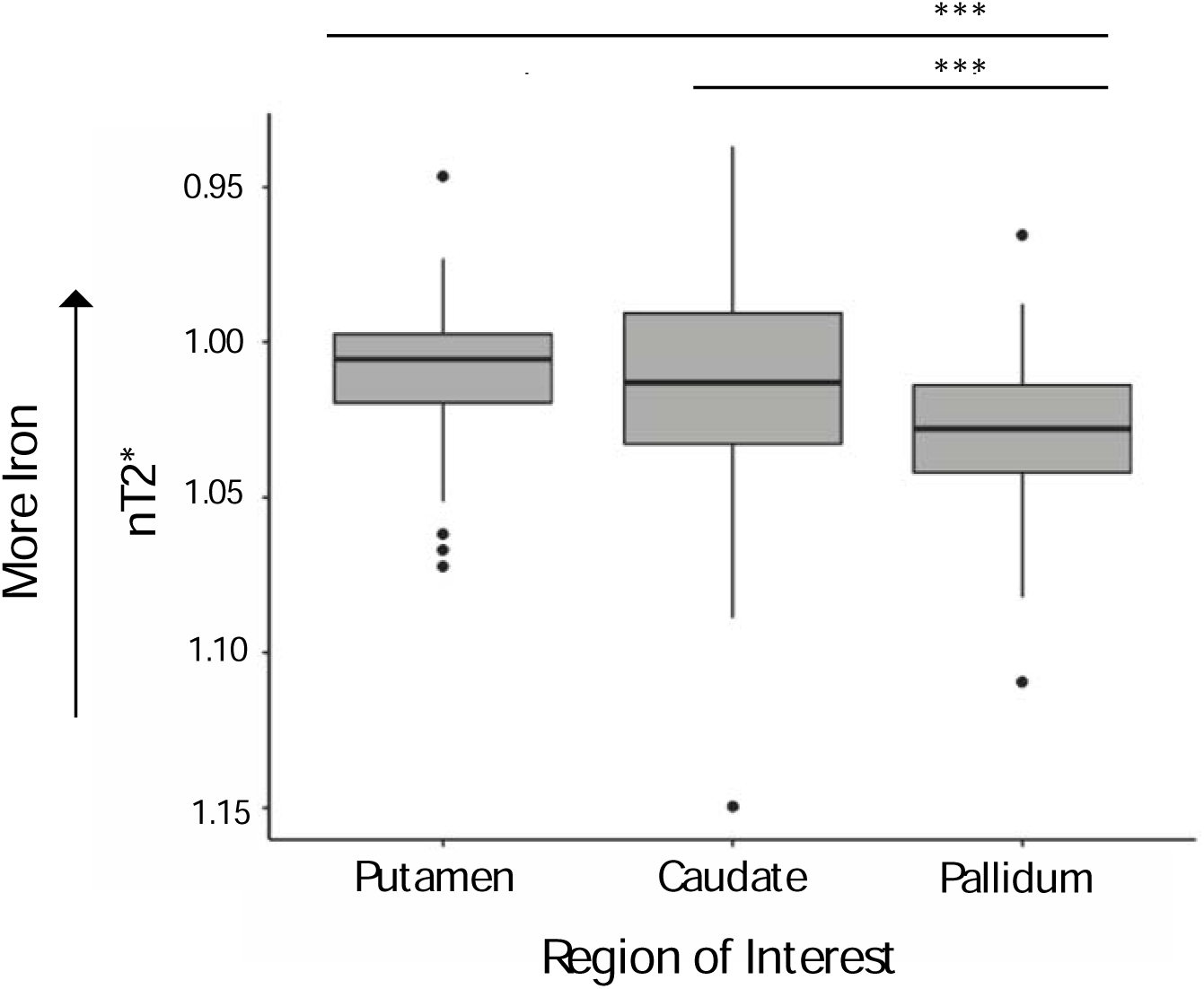
Boxplots that that show the distribution of nT2* for the three infant striatal regions. Lower values of nT2* are associated with greater iron levels. The pallidum had a nT2* signal that was significantly lower than the nT2* values in the caudate and putamen. Reliable differences were not found between the putamen and caudate.

### 3.2 Gestational and Postnatal Age Associations in Infancy

Linear models were used to assess the relationship between nT2* (indirectly related to iron) and age. A linear model that included gestational age at birth and postnatal age at scan as separate terms, while covarying for motion, sex, obstetric risk, birthweight, and maternal education, was run for each ROI (nT2* ∼ gestational age at birth (weeks) + postnatal age at scan (weeks) + mean framewise displacement + sex + obstetric risk + birth weight + maternal education). Motion was calculated from the fMRI time course, and mean framewise displacement (FD) was calculated for each participant. A distribution of FD values for all participants (M= 0.275) can be found in **Supplementary Figure 3**.

To assess if gestational age and postnatal age were colinear and impacting our regression models, we calculated the variance inflation factor and tolerance. When predictors were restricted to only the age terms, VIF was 1.07 and the tolerance was 0.93. Estimates of VIF remained low (below 1.2) when the remaining variables were estimated. Thus, collinearity is unlikely to have a meaningful impact on the results. Furthermore, in addition to the linear models described above, we repeated models considering gestational age and postnatal age in isolation as separate models. The results are similar to what was found below, and regression tables for those results are available in **Supplementary Table 1.**

#### 3.2.1 Pallidum

Gestational age was not a significant predictor of striatal nT2* (B= -2.530e-03, *p*=0.185). A linear model showed decreases in nT2* with postnatal age, reflecting indirect potential increases in tissue iron with age (B=-7.539e-03, p<0.001). There was no significant relationship between nT2* and motion (B=-6.860e-03, *p*=0.412). There was no relationship between obstetric risk (B=-4.507e-03, *p*=0.389) and nT2*, but there was a relationship with sex, with males having lower nT2* levels than females (B=-9.816e-03, *p*<0.05). There was also a significant association with birthweight and nT2* signal, with higher birthweights being associated with higher nT2* (B=2.123e-05, *p*<0.001). There was not a significant relationship between maternal education and nT2* signal (B=-3.160e-03, *p*=0.171). **Figure 2 A)** plots nT2* values with both gestational and postnatal age and **Supplementary Table 2** contains a regression table.

**Figure 2.**
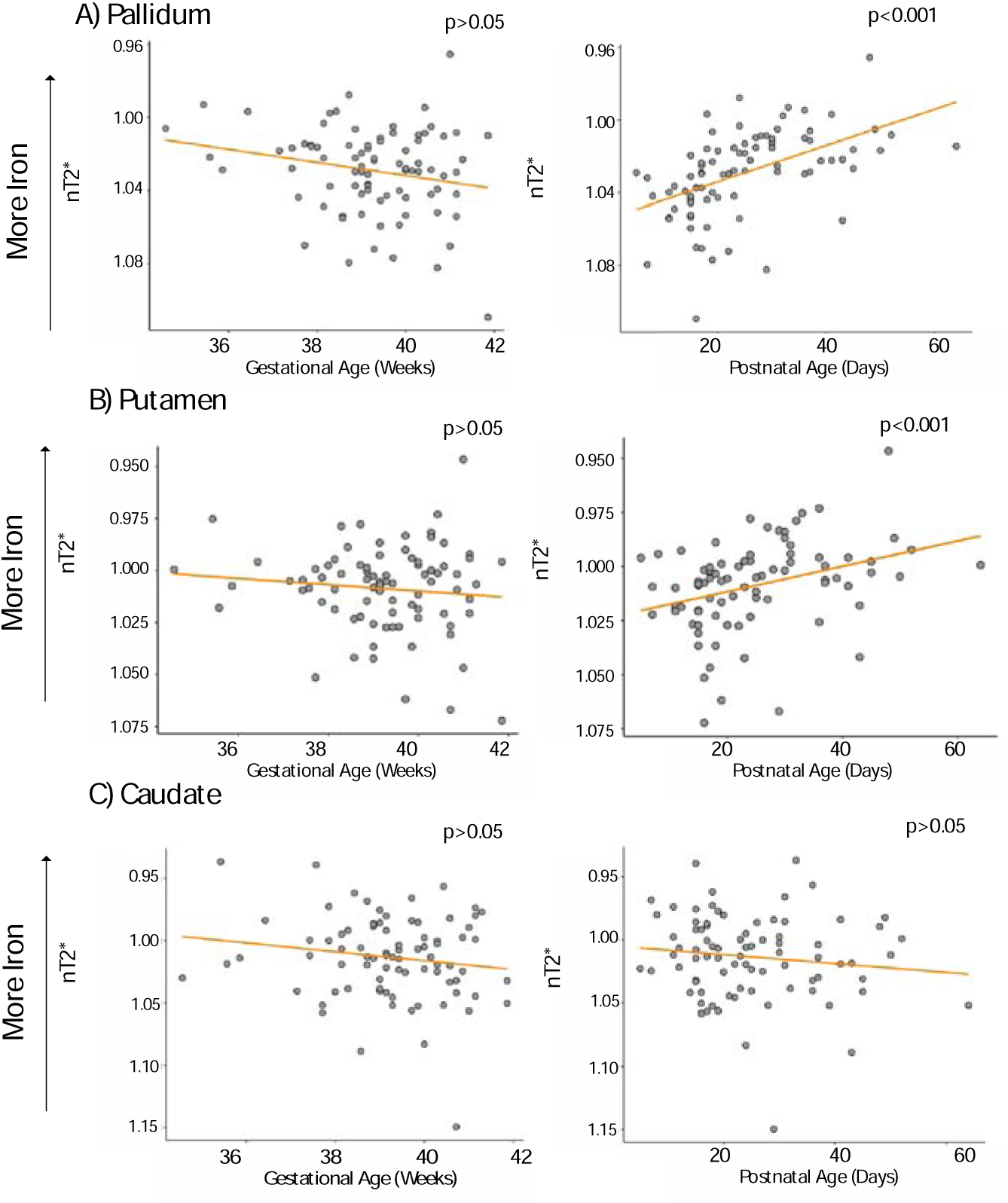
Gestational age (weeks) and postnatal age (days) plotted with nT2*, which we infer to indirectly measure iron deposition. Orange lines represent linear lines of best fit. We have inverted the y axis on all plots to reflect that iron is indirectly related to the inverse of nT2*.

#### 3.2.2 Putamen

Corresponding with the result seen in the pallidum, gestational age was not related to the nT2* signal (B=-2.943e-03, *p*=0.104), while there was a significant association with postnatal age (B=-5.091e-03, *p*<0.001), which we infer to reflect iron increasing with postnatal age. nT2* is plotted with both postnatal and gestational age in **Figure 2 B)**. Motion was not related to nT2* (B*=*-5.867e-03, *p*= 0.46). Like in the pallidum, there was a significant relationship between birthweight and nT2* signal (B=1.787e-05, *p*<0.001), where infants with higher birthweights had higher nT2*, which we infer to mean lower iron. There was a significant relationship between sex and nT2*, where males had lower nT2* than females (B=-1.317e-02, *p*<0.01), while there was no relationship between nT2* and obstetric risk (B=-4.557e-03, *p=*0.36). Maternal education was not related to nT2* (B=-3.370e-03, *p=*0.124). **Supplementary Table 2** contains a regression table with the full results.

After this, significant postnatal increases were observed in the pallidum (A) and the putamen (B), while gestational age was not related to nT2*. In the caudate (C) there was no relationship between age and nT2*.

#### 3.2.3 Caudate

Again, linear models assessed the relationship between age and developing nT2* signal. In contrast with the pallidum and putamen, there was not a significant association between postnatal age and nT2* (B=2.513e-03, *p*=0.249), and, consistent with the relationships found for the other two regions, there was not a relationship between gestational age and nT2* signal (B=- 1.624e-03, *p*=0.60). The relationship between age and nT2* signal is plotted in **Figure 2 C).**

Head motion (B=-1.358e-02, p= 0. 0.320) and sex (B=-1.374e-03, p=0.849) were not related to normalized T2* signal. Similarly with the pallidum and putamen, there was an association between nT2* and birthweight (B=2.829e-05, p<0.01), again where higher birthweights were associated with higher nT2* values. There was a relationship between obstetric risk and nT2* (B=-1.875e-02, p<0.05). Maternal education was not significantly related to nT2*(B=-5.011e-05, *p*=0.990). **Supplementary Table 2** contains a regression table with the full results.

A visualization of infant nT2* maps can be found in **Figure 3 A)**, which contains a mean voxelwise nT2* map for all participants with striatal regions of interest overlaid.

**Figure 3.**
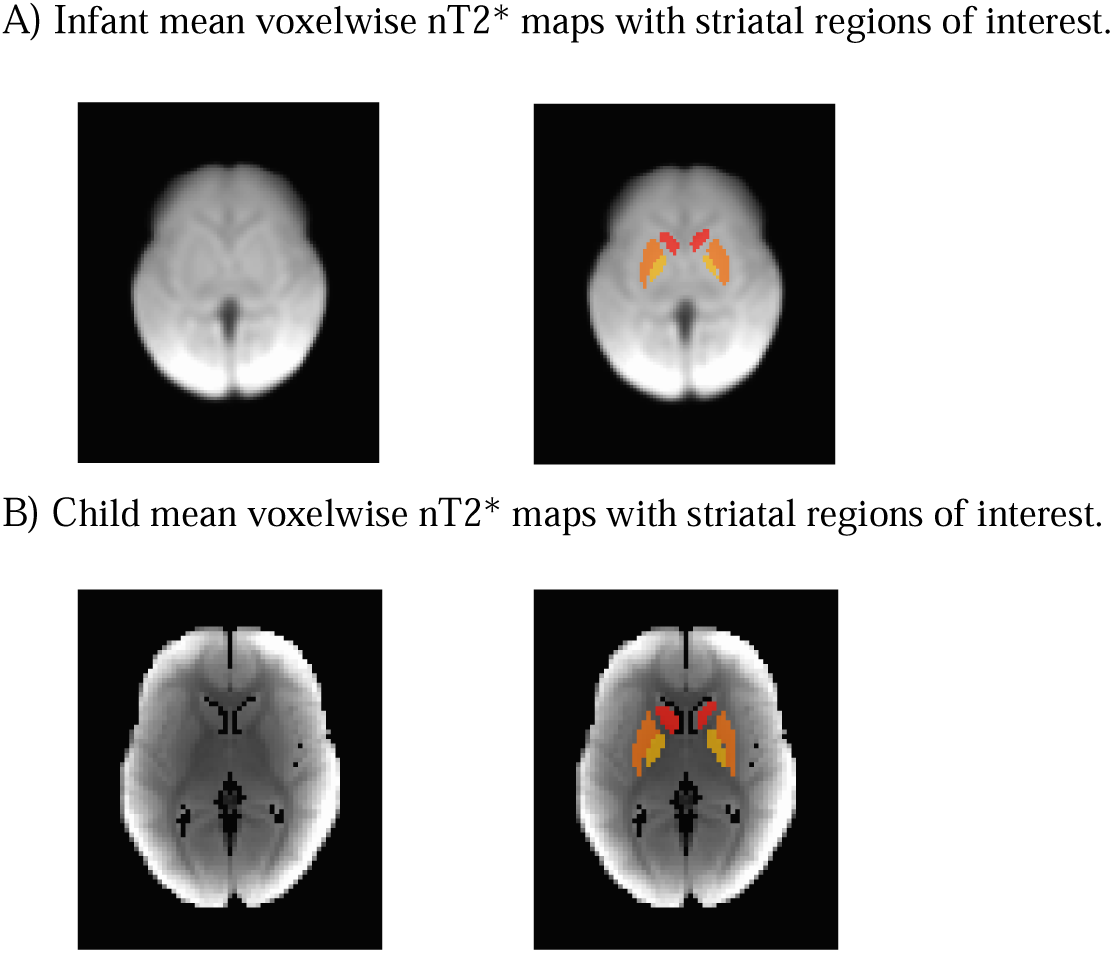
Group mean voxelwise nT2* maps for the N=83 infant (A) and N=26 child (B) participants. Scans are not harmonized or registered to the same template, and values cannot be directly compared between groups. However, the pattern of deposition between regions can be. The pallidum is darker, indicating lower nT2* and higher tissue iron, relative to the other regions, in children. This pattern is not seen in infants, where the pallidum has high nT2*. This indicates a shift between regional deposition between groups. Regions of interest include the caudate (red), putamen (orange), and pallidum (yellow).

### 3.3 Interaction Terms

As a post hoc test, we examined the interaction between gestational age and postnatal age. After mean centering, we found a significant interaction in the pallidum (B= - 2.001e-03, p<0.05) and putamen (B=-1.964e-03, p<0.05), with infants with shorter gestational ages changing less over time than those with longer gestational ages. We did not find a significant relationship in the caudate (B=-2.540e-03, p>0.05). These results are plotted in **Supplementary Figure 4**. As we did not have an a priori hypothesis for the interaction, we caution against overinterpreting the results

### 3.4 Relationship between birthweight and brain volume

It is possible that infants who have larger birthweights also have larger brain volumes. An association between birthweight and brain volume might mean the relationship between birthweight and nT2* was being partially driven by brain volume, representing a confounding variable. To test this, we used segmentations from the structural images to test if volume, excluding the ventricles, was related to birth weight. There was a small correlation between brain volume and birth weight (r(81)= 0.23, p=0.037). However, there was not a relationship between the number of non-zero voxels in a subject’s nT2* image and birthweight, (r(81)= -.13, p=0.258). To validate that the effect of birthweight was present after adjusting for brain volume, we also added brain volume, as calculated from the structural images, to our model for the pallidum, our region with the largest age effect. We found that the relationship between nT2* and postnatal age (B=-3.866e-03, p<0.05) continued to persist even when characterizing brain volume. Increased birthweight (B=2.149e-05, p<0.001) was associated with higher nT2* and increased brain volume (B=-1.996e-07, p<0.01 was associated with lower nT2*.

### 3.5 Distribution of tissue iron at 5 years

To examine how the distribution of tissue iron deposition shifts in early childhood, we completed an exploratory analysis in 26 children that were scanned as neonates. The data presented here is not harmonized with the infant data, and values should not be directly compared between groups. However, differences in confounds, such as scanners and sequences, likely affect all striatal regions equally within each age group. Thus, it is possible to compare the pattern of iron deposited between regions in the adults to the pattern of iron deposited between regions in the infants. In contrast to the infant tissue iron data, the pallidum (M=0.670, SD=0.049) was found to have the lowest nT2* (indirectly reflecting high iron), while the putamen (M=0.815, SD=0.045) and caudate had lower values (M=0.817, SD=0.043). There was a significant difference in nT2* between the pallidum and putamen (t(25)=22.29, p<0.001), a pattern that was also found for the pallidum and caudate (t(25)=16.60, p<0.001). The putamen and caudate were not significantly different from each other (t(25)=-049, p=0.627). The distribution of tissue iron among the three regions is plotted in **Figure 4**. A visualization of child nT2* maps, as a mean voxelwise image with striatal regions of interest overlaid, can be found in **Figure 3 B)**.

**Figure 4.**
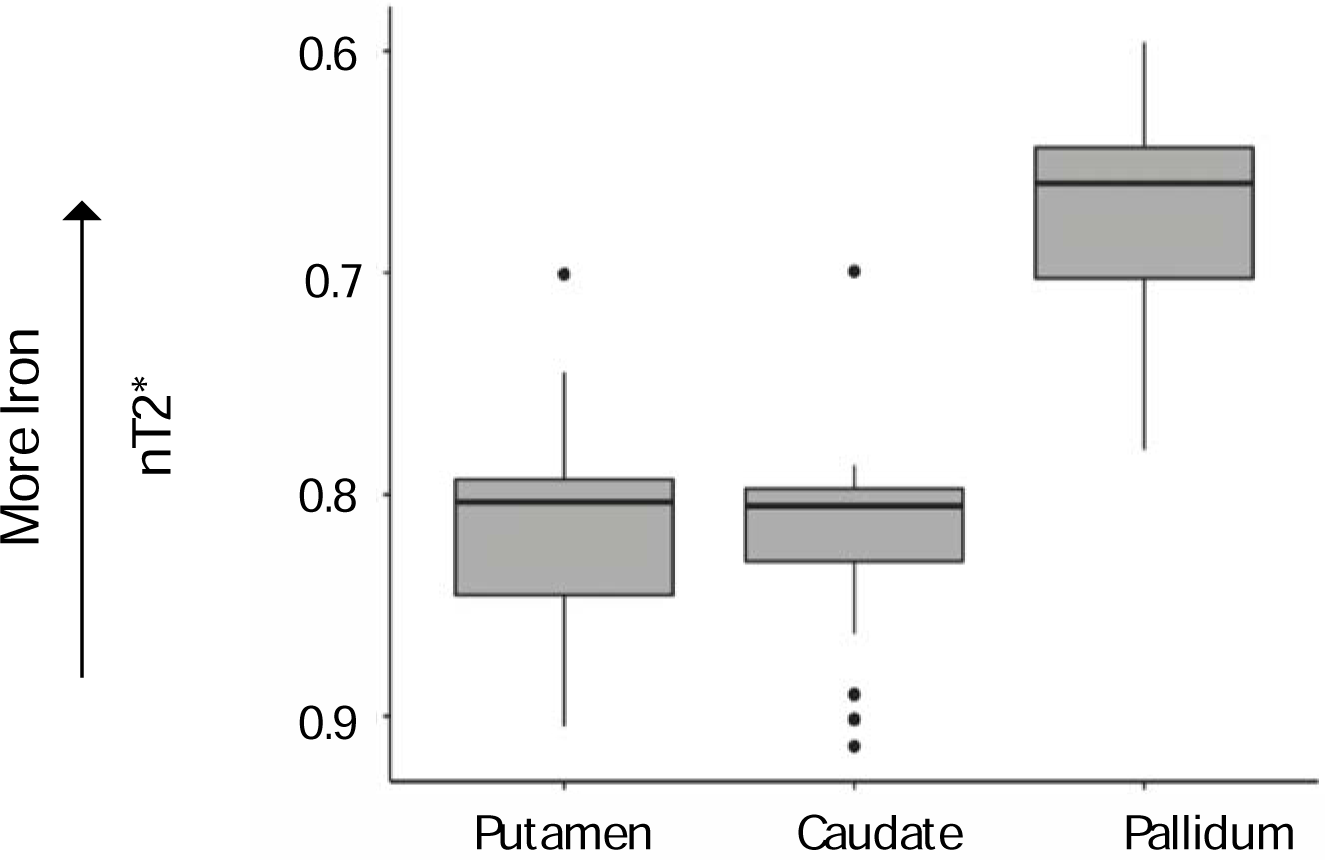
Boxplots depicting the distribution of nT2* in the striatum in preschool children. Lower values of nT2* are linked to higher iron for the N=26 children that were scanned neonatally. The pallidum had significantly greater iron than the putamen and the caudate.

Iron increased significantly with postnatal age in the pallidum (B=-0.023838, p<0.05), but not in the putamen (B=0.0035174, p=0.733) or caudate (B= 0.009266, p=0.343). There was no relationship with gestational age in the pallidum (B=-0.002806, p=0.710), putamen (B=0.0002552, p=0.974), or caudate (B=0.003645, p=0.620). The relationship between postnatal age and nT2* is plotted in **Supplementary Figure 5** and a regression table is in **Supplementary Table 3.**

## 4.0 Discussion

For the first time, we identify postnatal age associations for striatal subregions in early life, while accounting for gestational age. If gestation had been an extension of what we found postnatally, we would have expected linear increases for both variables. Instead, this result exemplifies the postnatal period as one of rapid change, while controlling for gestational age. The pallidum and putamen had significant postnatal indirect increases in iron deposition with age, but there were no significant increases with gestational age. In contrast, the caudate did not have a significant relationship with either postnatal or gestational age.

Differences across regions may reflect different maturational timelines in the development of striatal neurophysiology. When considering the overall level of nT2* for the three regions, the putamen and caudate had comparable levels of in infancy, while the pallidum’s nT2* has the potential to reflect significantly lower iron deposition. To examine how the distribution among subregions shifts in early life, we conducted an initial exploratory analysis, using a small longitudinal sample (N=26) of infants who were followed up and scanned at preschool age. As the infant and preschool scans occurred on different scanners with different sequences, direct comparisons between the infant and preschool nT2* values should not be made. However, any confounding variables likely affect striatal subregions equally within groups, and the relative differences between regions at each timepoint can be used to illustrate trajectories of developmental change. At preschool age, similarly to the infant data, the caudate and putamen did not have significantly different nT2*. In contrast to the infant sample, we can indirectly infer the pallidum had significantly more iron accrual than the caudate and putamen. Our inferences in childhood are consistent with data from large cohorts in adolescence and adults ^2, 16^, where the pallidum is the most iron rich region and in studies with children, where the pallidum can be visually delineated from other striatal subregions (Zhang et al (2019)). Taken together, these results demonstrate early childhood as a potential time of critical maturation for the pallidum, as it moves from having the highest nT2* to a pattern that is adult-like. Our results are in line with early work demonstrating whole striatum and regional postnatal increases in iron deposition in infancy. Interestingly, our work corresponds with Ning et al (2019), who found age-related change earlier in the pallidum and putamen than the caudate.

The rapid age-related change exhibited by the pallidum in early postnatal life could reflect early motor development. The pallidum plays a large role in motor function because of its widespread connections to all the basal ganglia nuclei and to many sensorimotor areas ^42^. Additionally, the pallidum plays a role in stopping actions, which also may be more relevant in postnatal life ^43^. The ventral pallidum has also been proposed to play a role in reward and motivation ^44^. It’s possible that the postnatal change seen here could reflect a combination of both the maturation of motor and reward systems. Early motor development could initiate initial postnatal change, then, as reward systems come rapidly online, this could help to drive the eventual increased iron deposition over other regions. Motor development also has the potential to drive iron deposition in the putamen through its strong connectivity to both the primary motor cortex and the supplementary motor area, where stimulating those regions, in animals and humans, results in neuronal firing in the putamen ^45–47^. The putamen plays a role in motor initiation and reversal learning ^48, 49^, which may particularly be relevant in the first days and months of postnatal life as infants have the space and opportunity to perform more complex and voluntary movements.

In contrast, the caudate had low nT2* but did not significantly increase its iron content in the postnatal period. These results are in line with fetal work measuring relative caudate volume, which showed less age-related change at later gestational ages ^50^. When looking at striatal iron in childhood, iron in the caudate is often related to cognitive abilities ^16, 34, 35^. It has been proposed that the caudate is involved in both the planning and the execution of goal-oriented outcomes or strategies ^51–54^. The caudate has substantial connections to prefrontal regions ^55, 56^, and there is evidence that frontal areas are already functioning with substantial maturity in early perinatal life ^57, 58^, although with less reproducibility than primary sensory networks ^59^. Initial iron deposition could be driven by this. Maturation may then slow in comparison with other brain other areas that are experiencing rapid myelination and gyrification and then again may become evident as the trajectory of brain growth stabilizes in the second half of the first year. Thus, this would reflect overall developmental processes of sensorimotor systems maturing before cognitive processes ^60^. Alternatively, the tail of the caudate is hypothesized to be involved in visual processing^61^, which matures early^62, 63^, and could be driving initial iron deposition.

When examining potentially confounding factors, sex was related to iron deposition in the putamen and pallidum, where males had higher iron deposition than females. Although this relationship is inconsistent across regions, and some studies have failed to find sex differences in early male and female brain development ^64–66^, there are some reliable differences between male and female brains in infancy. For example, male brains are often larger than female brains ^67^, and there is some evidence that the putamen is larger in males than females early in childhood ^68, 69^. Future work with larger samples and multiple imaging modalities, such as iron and brain volume, should continue to explore this relationship.

We found a significant relationship between nT2* and birthweight for all the three regions in a sample not enriched for preterm birth or iron deficiency, where higher birthweight was associated with increased nT2*. Birthweight was not significantly related to motion or total brain volume in the participants, so it is unlikely that larger infants moved more in the scanner or brain volume was a confounding measure. Gestational age is related to birthweight, and infants with longer gestational ages have higher birthweights, but the association with birth weight persisted after controlling for both gestational and postnatal age ^70^. Infant birth weight is related to many maternal characteristics, including the weight of the mother and iron levels. In a retrospective study of 11,581 Chinese infants, lower maternal serum ferratin in the third trimester was related to higher birthweights after adjusting for gestational age ^71^. Future work should seek to understand this association and should collect blood iron levels from neonates at birth and at the time of scan to characterize the relationship more fully.

Our study was designed to characterize normative development, and the infants were born roughly at term equivalent age. We did not observe increases in nT2* that were related to gestational age, suggesting that significant striatal maturation takes place after birth perhaps requiring by external environmental stimuli. However, a larger range of gestational ages, including preterm infants, may reveal possible associations that we may have missed. Future work should seek to examine tissue iron deposition for a range of gestational ages to fully characterize whether increases in iron are associated with increases in gestational age. However, we would like to note that this cohort has roughly equivalent variation in GA as in postnatal age at scan. Thus, we have an appropriate amount of variation to compare between age variables.

In comparison with previous work, our study used a larger sample size (N=83) of primarily younger infants. However, we do not have enough participants to separate out exclusively perinatal development from later neonatal development, and future work with larger samples should seek to characterize whether change is greatest around the time of birth or in infancy. Finally, our study used a method of indirectly measuring tissue iron that has been successfully used in adolescence and in adulthood ^2, 17^ and, here, for the first time used in infancy. The strength of this method is that it can be used with existing resting state images without additional sequences or specific acquisition parameters, allowing for large samples of infant participants and decreased scan time. In adolescence and adults, using T2* from rsfMRI to estimate iron shows close correspondence with other MRI sequences that are established as markers of iron, including R2* ^17^. However, the T2* signal is also sensitive to changes in water content, and the water content, along with myelin and tissue density, shifts rapidly within the brain in early life. The brain is 88% water at term and this decreases to approximately 85% 6 months after birth ^72, 73^. To ensure the developmental decreases in nT2* observed in the striatum are not specific to general shifts in water concentration, calculating normalized T2* involves normalizing each voxel to the whole brain median, benchmarking a subject’s nT2* map against their overall development and shifts in water. To further substantiate this, future work should seek to compare our method with concurrent R2* imaging in the same infants to confirm what has already been established in adolescence and in later childhood, close correspondence between the two imaging acquisitions.

The relationship between brain iron and blood iron remains a topic for future research, as we did not screen for iron deficiency in our normative sample. It is currently unknown what the impact of maternal or infant blood iron is on concurrent brain iron deposition. Future work should seek to obtain blood samples from the mother during pregnancy, from the infant at birth, and then from the infant at the time of the MRI scan to begin to model these factors and determine if increased iron availability leads to increased brain tissue iron deposition in normative samples, and in clinical populations whether hypoxia and anemia are related to increased deposition. Hypoxia also has the potential to affect the T2* signal because deoxyhemoglobin, like iron, is paramagnetic. Despite this, deoxyhemoglobin is unlikely to impact nT2*. nT2* is a relative measurement, and as deoxyhemoglobin concentration is likely similar in the striatum as in the whole brain, any affect would likely be removed through normalization. Furthermore, longitudinal fetal imaging would allow for an additional brain timepoint and may help to characterize the relationship between gestational age, brain iron, and later clinical characteristics, such as anemia.

Finally, this work informs both normative and abnormal development, such as in mental illness. Mapping periods of sensitive change that may be specific to motor, cognitive, and reward systems allow us to relate these early measures to more sophisticated neurodevelopmental tests in later childhood. The trajectory of and associations between brain iron and behavior may provide additional predicative predictive utility for clinical models trying to characterize abnormal development. These models may be especially useful for related iron-related disorders, such as anemia, that occur during key early developmental windows and where interventions may need timely intervention to bring brain development back in line with typically developing peers. This work is a first step in using nT2* to characterize brain tissue iron in early development. This method has the potential to be widely used in existing datasets, and future work linking the results here to the many outstanding questions offers many interesting applications in both clinical disorders and normative development.

## Declarations

Damien A. Fair is a patent holder on the Framewise Integrated Real-Time Motion Monitoring (FIRMM) software. He is also a co-founder of Turing Medical Inc. The nature of this financial interest and the design of the study have been reviewed by two committees at the University of Minnesota. They have put in place a plan to help ensure that this research study is not affected by the financial interest.

## Supplementary Figures

**Supplementary Figure 1.**
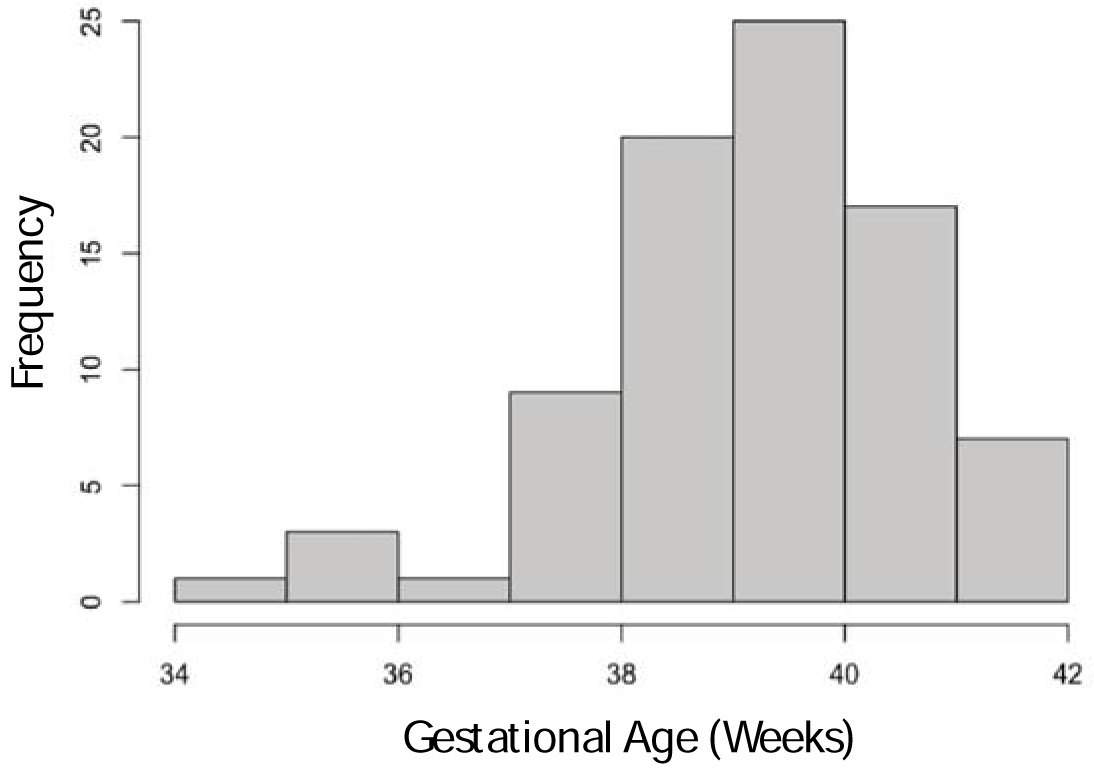
The distribution of gestational age for the 83 participants was included in the statistical models.

**Supplementary Figure 2.**
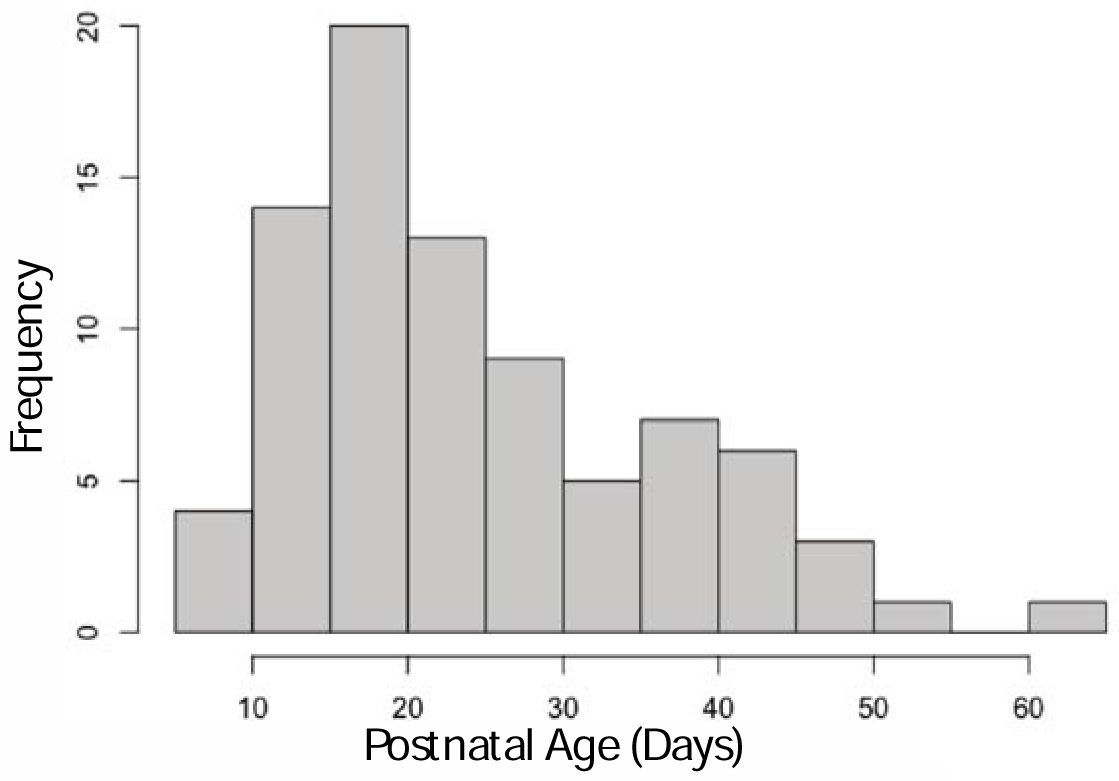
The distribution of postnatal age for the 83 participants was included in the statistical models.

**Supplementary Figure 3.**
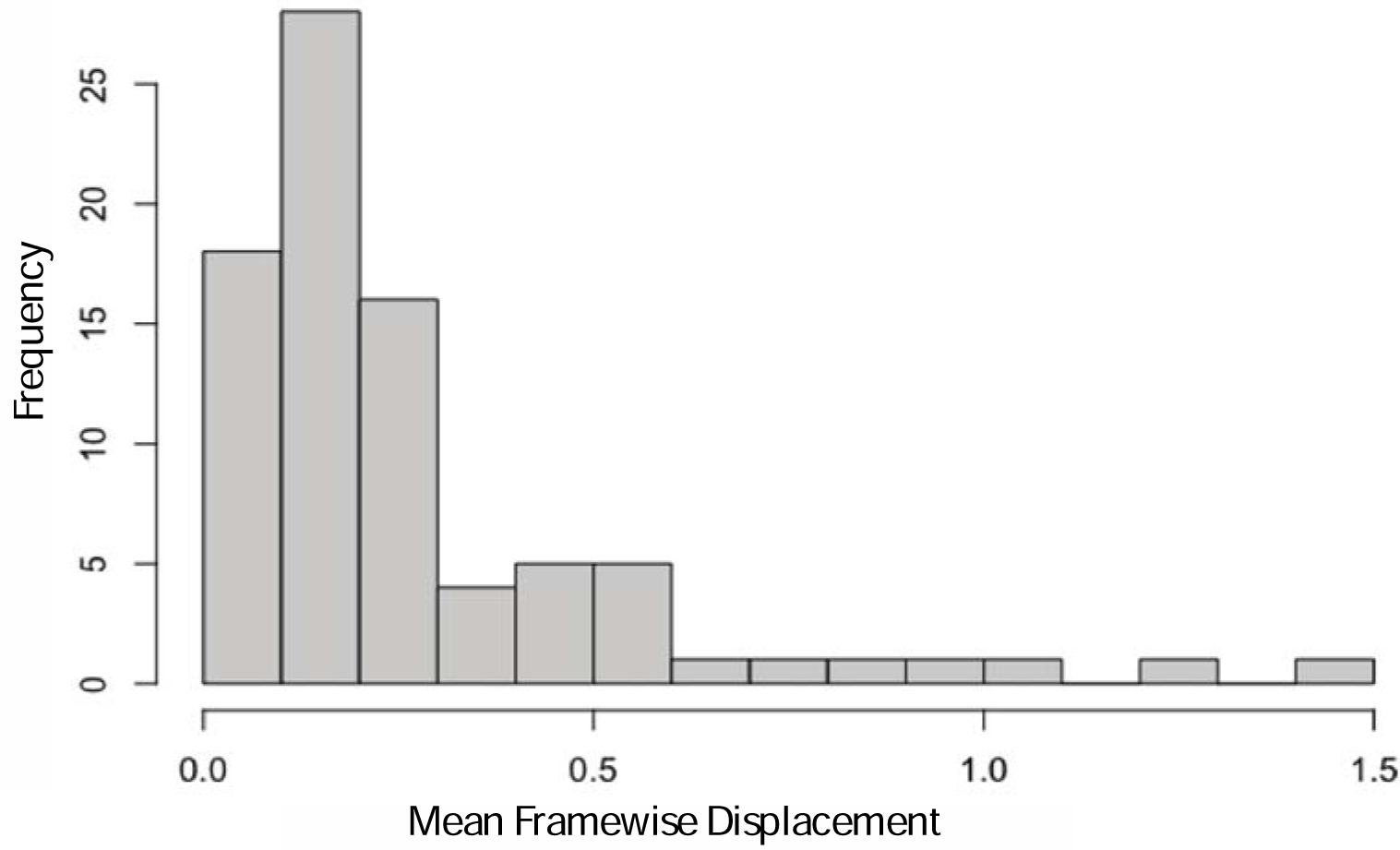
The distribution of mean framewise displacement for the 83 participants was included in the statistical models.

**Supplementary Table 1.**
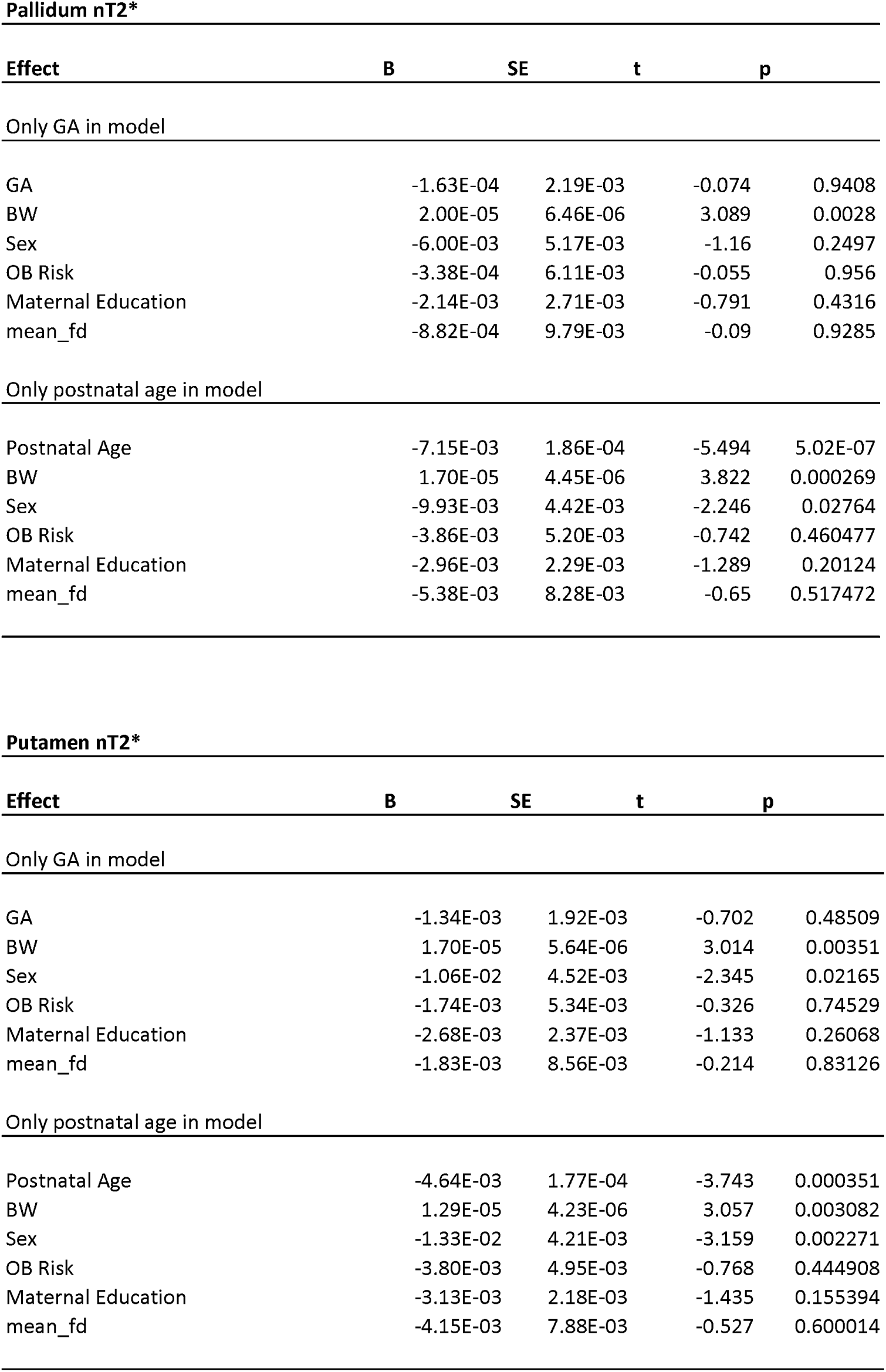
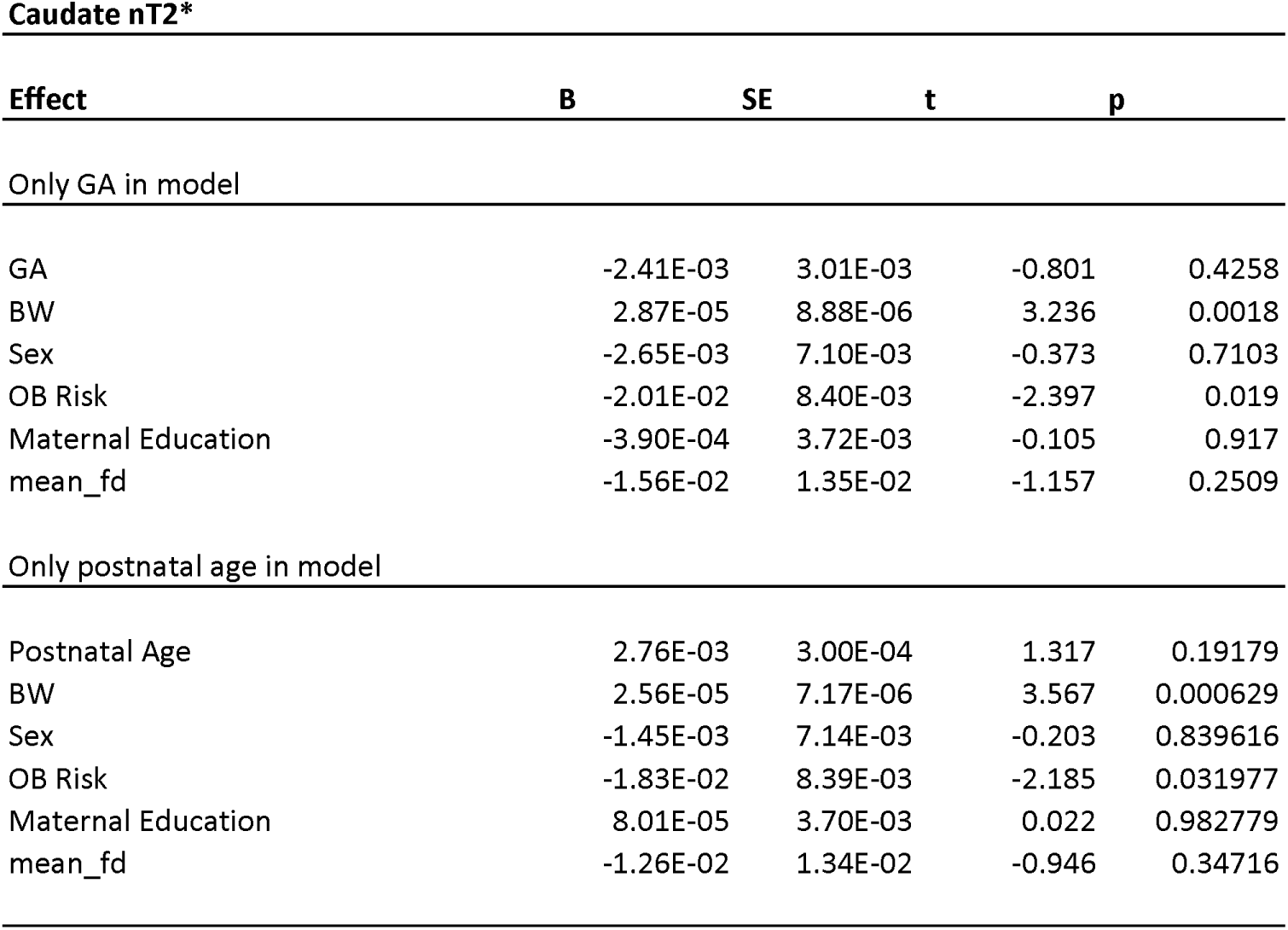
Regression tables for the pallidum, putamen and caudate with only one age term per model. Results are similar to what was found when both age terms were included. The nT2* in the pallidum and putamen both had an assocation with postnatal age but not with gestational age. There were not any age-related associations in the caudate.

| Effect | B | SE | t | p |
| --- | --- | --- | --- | --- |
| Only GA in model |  |  |  |  |
| GA | -1.63E-04 | 2.19E-03 | -0.074 | 0.9408 |
| BW | 2.00E-05 | 6.46E-06 | 3.089 | 0.0028 |
| Sex | -6.00E-03 | 5.17E-03 | -1.16 | 0.2497 |
| OB Risk | -3.38E-04 | 6.11E-03 | -0.055 | 0.956 |
| Maternal Education | -2.14E-03 | 2.71E-03 | -0.791 | 0.4316 |
| mean_fd | -8.82E-04 | 9.79E-03 | -0.09 | 0.9285 |
| Only postnatal age in model |  |  |  |  |
| Postnatal Age | -7.15E-03 | 1.86E-04 | -5.494 | 5.02E-07 |
| BW | 1.70E-05 | 4.45E-06 | 3.822 | 0.000269 |
| Sex | -9.93E-03 | 4.42E-03 | -2.246 | 0.02764 |
| OB Risk | -3.86E-03 | 5.20E-03 | -0.742 | 0.460477 |
| Maternal Education | -2.96E-03 | 2.29E-03 | -1.289 | 0.20124 |
| mean_fd | -5.38E-03 | 8.28E-03 | -0.65 | 0.517472 |

**Putamen nT2\***
| Effect | B | SE | t | p |
| --- | --- | --- | --- | --- |
| Only GA in model |  |  |  |  |
| GA | -1.34E-03 | 1.92E-03 | -0.702 | 0.48509 |
| BW | 1.70E-05 | 5.64E-06 | 3.014 | 0.00351 |
| Sex | -1.06E-02 | 4.52E-03 | -2.345 | 0.02165 |
| OB Risk | -1.74E-03 | 5.34E-03 | -0.326 | 0.74529 |
| Maternal Education | -2.68E-03 | 2.37E-03 | -1.133 | 0.26068 |
| mean_fd | -1.83E-03 | 8.56E-03 | -0.214 | 0.83126 |
| Only postnatal age in model |  |  |  |  |
| Postnatal Age | -4.64E-03 | 1.77E-04 | -3.743 | 0.000351 |
| BW | 1.29E-05 | 4.23E-06 | 3.057 | 0.003082 |
| Sex | -1.33E-02 | 4.21E-03 | -3.159 | 0.002271 |
| OB Risk | -3.80E-03 | 4.95E-03 | -0.768 | 0.444908 |
| Maternal Education | -3.13E-03 | 2.18E-03 | -1.435 | 0.155394 |
| mean_fd | -4.15E-03 | 7.88E-03 | -0.527 | 0.600014 |

*Supplementary Table 1.* Regression tables for the pallidum, putamen and caudate with only one age term per model.
| **Caudate nT2\*** |  |  |  |  |
| --- | --- | --- | --- | --- |
| **Effect** | **B** | **SE** | **t** | **p** |
| Only GA in model |  |  |  |  |
| GA | -2.41E-03 | 3.01E-03 | -0.801 | 0.4258 |
| BW | 2.87E-05 | 8.88E-06 | 3.236 | 0.0018 |
| Sex | -2.65E-03 | 7.10E-03 | -0.373 | 0.7103 |
| OB Risk | -2.01E-02 | 8.40E-03 | -2.397 | 0.019 |
| Maternal Education | -3.90E-04 | 3.72E-03 | -0.105 | 0.917 |
| mean_fd | -1.56E-02 | 1.35E-02 | -1.157 | 0.2509 |
| Only postnatal age in model |  |  |  |  |
| Postnatal Age | 2.76E-03 | 3.00E-04 | 1.317 | 0.19179 |
| BW | 2.56E-05 | 7.17E-06 | 3.567 | 0.000629 |
| Sex | -1.45E-03 | 7.14E-03 | -0.203 | 0.839616 |
| OB Risk | -1.83E-02 | 8.39E-03 | -2.185 | 0.031977 |
| Maternal Education | 8.01E-05 | 3.70E-03 | 0.022 | 0.982779 |
| mean_fd | -1.26E-02 | 1.34E-02 | -0.946 | 0.34716 |

**Supplementary Table 2.** Regression tables for the linear models used in the infant pallidum, putamen and caudate. The pallidum and putamen had associations between nT2* and postnatal age (weeks) but not with gestational age. The caudate didn’t have an association between nT2* and either of the age terms.

| **Pallidum nT2\*** |  |  |  |  |
| --- | --- | --- | --- | --- |
| **Effect** | **B** | **SE** | **t** | **p** |
| Postnatal Age | -7.54E-03 | 1.33E-03 | -5.682 | 2.41E-07 |
| GA | -2.53E-03 | 1.89E-03 | -1.338 | 0.185047 |
| BW | 2.12E-05 | 5.44E-06 | 3.902 | 0.000207 |
| Sex | -9.82E-03 | 4.40E-03 | -2.23 | 0.028724 |
| OB Risk | -4.51E-03 | 5.20E-03 | -0.867 | 0.388667 |
| Maternal Education | -3.16E-03 | 2.29E-03 | -1.382 | 0.171142 |
| mean_fd | -6.86E-03 | 8.31E-03 | -0.825 | 0.411752 |
| **Putamen nT2\*** |  |  |  |  |
| **Effect** | **B** | **SE** | **t** | **p** |
| Postnatal Age | -5.09E-03 | 1.26E-03 | -4.054 | 0.000122 |
| GA | -2.94E-03 | 1.79E-03 | -1.643 | 0.104474 |
| BW | 1.79E-05 | 5.15E-06 | 3.47 | 0.000866 |
| Sex | -1.32E-02 | 4.17E-03 | -3.161 | 0.002268 |
| OB Risk | -4.56E-03 | 4.92E-03 | -0.926 | 0.357292 |
| Maternal Education | -3.37E-03 | 2.17E-03 | -1.557 | 0.123726 |
| mean_fd | -5.87E-03 | 7.87E-03 | -0.746 | 0.458073 |
| **Caudate nT2\*** |  |  |  |  |
| **Effect** | **B** | **SE** | **t** | **p** |
| Postnatal Age | 2.51E-03 | 2.16E-03 | 1.163 | 0.2487 |
| GA | -1.62E-03 | 3.08E-03 | -0.527 | 0.59991 |
| BW | 2.83E-05 | 8.86E-06 | 3.192 | 0.00206 |
| Sex | -1.37E-03 | 7.17E-03 | -0.192 | 0.84855 |
| OB Risk | -1.88E-02 | 8.47E-03 | -2.214 | 0.02986 |
| Maternal Education | -5.01E-05 | 3.73E-03 | -0.013 | 0.98931 |
| mean_fd | -1.36E-02 | 1.35E-02 | -1.003 | 0.31906 |

**Supplementary Figure 4.**
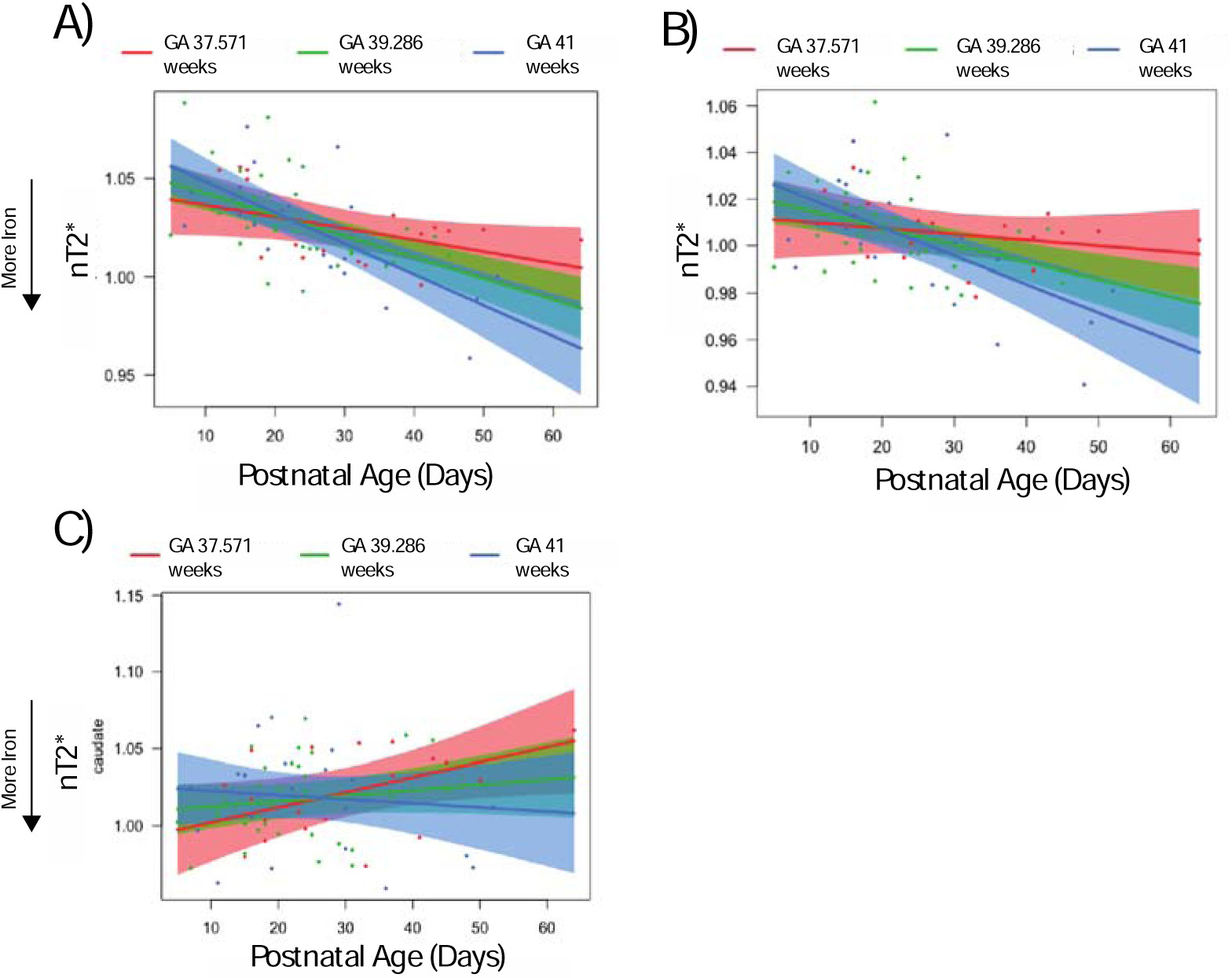
Visualization of the regression models, containing the interaction terms and covariates, divided into cross-sections. Cross-sections represent the 10^th^, 50^th^, and 90^th^ quantiles of gestational age, with postnatal age on the x axis. Points represent partial residuals, plotted with prediction lines and confidence bands. The pallidum is in A); the putamen is in B); the caudate is in C). Only the putamen and the pallidum had significant interactions, with infants with shorter gestational ages appearing to change less over time.

**Supplementary Figure 5.**
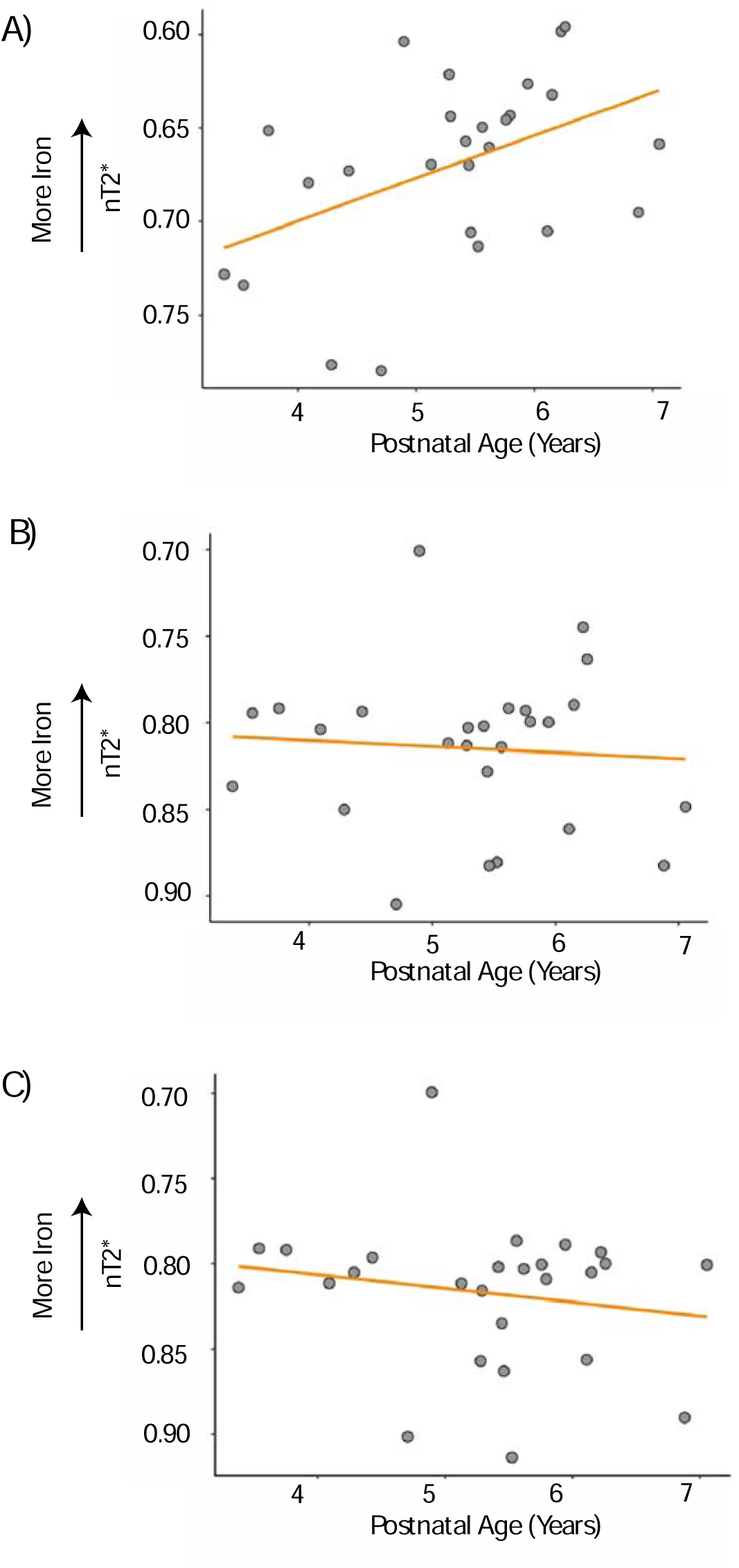
Postnatal age plotted with nT2* values for the infants scanned longitudinally longitudinally as preschoolers. Orange lines represent the lines of best fit. In the pallidum (A), postnatal age was significantly associated with iron deposition, but not for the putamen (B) or the caudate (C).

**Supplementary Table 3.**
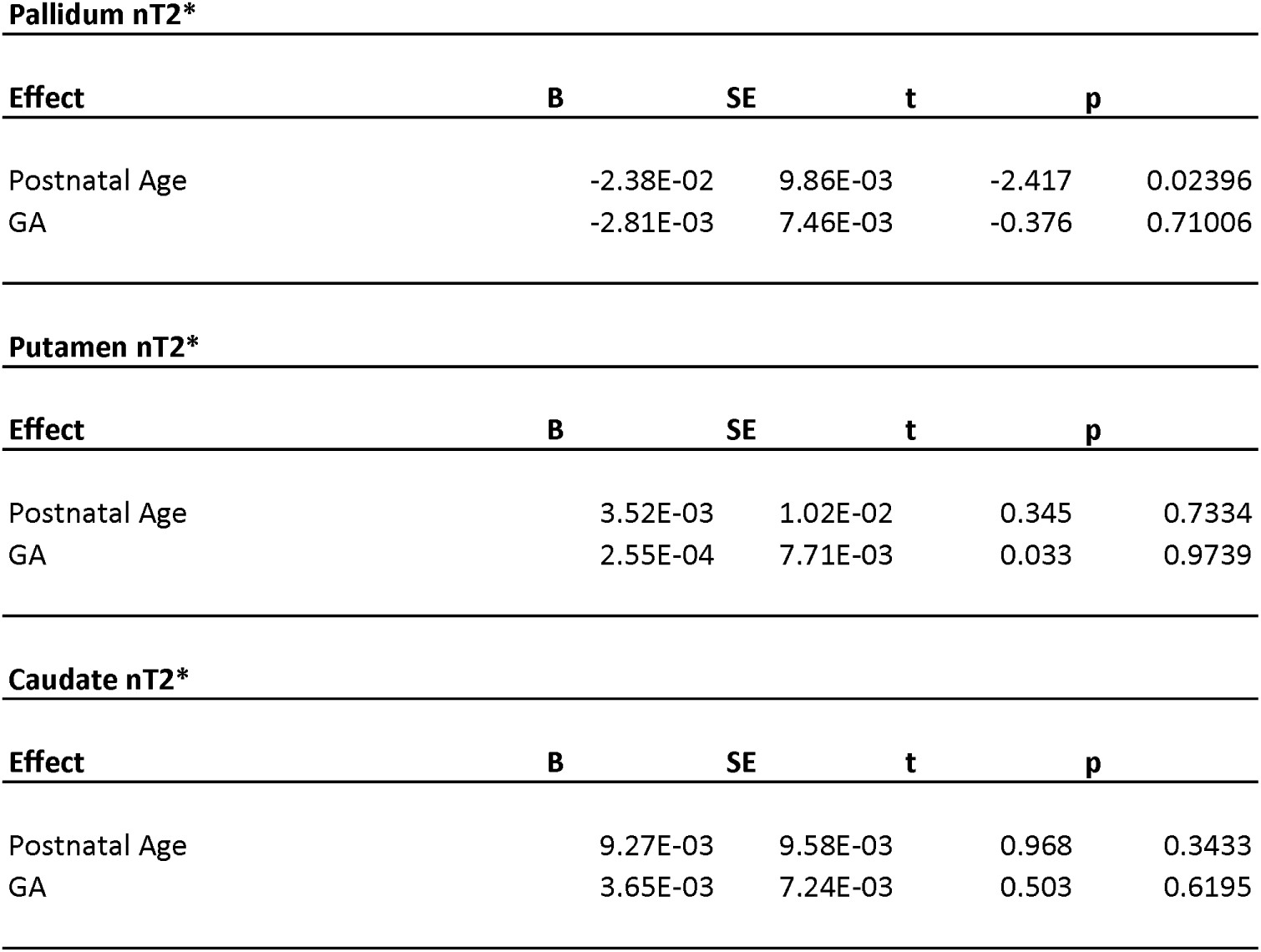
Regression table for the preschooler linear models. The pallidum had an association between postnatal age and nT2*, but there were no other significant associations with age.

## References

1. Shepherd, G. M. G. Corticostriatal connectivity and its role in disease. Nat. Rev. Neurosci. 14, 278–291 (2013).

2. Larsen, B. & Luna, B. In vivo evidence of neurophysiological maturation of the human adolescent striatum. Dev. Cogn. Neurosci. 12C, 74–85 (2015).

3. Thomas, L. O., Boyko, O. B., Anthony, D. C. & Burger, P. C. MR detection of brain iron. AJNR Am. J. Neuroradiol. 14, 1043–1048 (1993).

4 Parr, A. C., et al. Dopamine-related striatal neurophysiology is associated with specialization of frontostriatal reward circuitry through adolescence. bioRxiv 2020.06.24.169847 (2020) doi:10.1101/2020.06.24.169847.

5. Longitudinal Development of Brain Iron Is Linked to Cognition in Youth | Journal of Neuroscience. https://www.jneurosci.org/content/40/9/1810.

6. Brass, S. D., Chen, N., Mulkern, R. V. & Bakshi, R. Magnetic resonance imaging of iron deposition in neurological disorders. Top. Magn. Reson. Imaging TMRI 17, 31–40 (2006).

7. Hallgren, B. & Sourander, P. The effect of age on the non-haemin iron in the human brain. J. Neurochem. 3, 41–51 (1958).

8. Ward, R. J., Zucca, F. A., Duyn, J. H., Crichton, R. R. & Zecca, L. The role of iron in brain ageing and neurodegenerative disorders. Lancet Neurol. 13, 1045–1060 (2014).

9. Paul, B. T., Manz, D. H., Torti, F. M. & Torti, S. V. Mitochondria and Iron: current questions. Expert Rev. Hematol. 10, 65–79 (2017).

10. Horowitz, M. P. & Greenamyre, J. T. Mitochondrial Iron Metabolism and Its Role in Neurodegeneration. J. Alzheimers Dis. JAD 20, S551–S568 (2010).

11. Ortega, R., Cloetens, P., Devès, G., Carmona, A. & Bohic, S. Iron Storage within Dopamine Neurovesicles Revealed by Chemical Nano-Imaging. PLOS ONE 2, e925 (2007).

12. Youdim, M. B. H. & Green, A. R. Iron deficiency and neurotransmitter synthesis and function. Proc. Nutr. Soc. 37, 173–179 (1978).

13. Youdim, M. B. H. Monoamine oxidase inhibitors, and iron chelators in depressive illness and neurodegenerative diseases. J. Neural Transm. Vienna Austria 1996 125, 1719–1733 (2018).

14. Zhang, Y. et al. Longitudinal data for magnetic susceptibility of normative human brain development and aging over the lifespan. Data Brief 20, 623–631 (2018).

15. Li, W. et al. Differential developmental trajectories of magnetic susceptibility in human brain gray and white matter over the lifespan. Hum. Brain Mapp. 35, 2698–2713 (2014).

16. Larsen, B. et al. Longitudinal Development of Brain Iron Is Linked to Cognition in Youth. J. Neurosci. 40, 1810–1818 (2020).

17. Parr, A. C. et al. Contributions of dopamine-related basal ganglia neurophysiology to the developmental effects of incentives on inhibitory control. Dev. Cogn. Neurosci. 54, 101100 (2022).

18. Parr, A. C. et al. Dopamine-related striatal neurophysiology is associated with specialization of frontostriatal reward circuitry through adolescence. Prog. Neurobiol. 101997 (2021) doi:10.1016/j.pneurobio.2021.101997.

19. Monk, C. et al. Maternal prenatal iron status and tissue organization in the neonatal brain. Pediatr. Res. 79, 482–488 (2016).

20. Rao, R. & Georgieff, M. K. Perinatal aspects of iron metabolism. Acta Paediatr. Oslo Nor. 1992 Suppl. 91, 124–129 (2002).

21. Tamura, T. et al. Cord serum ferritin concentrations and mental and psychomotor development of children at five years of age. J. Pediatr. 140, 165–170 (2002).

22. Walter, T., De Andraca, I., Chadud, P. & Perales, C. G. Iron deficiency anemia: adverse effects on infant psychomotor development. Pediatrics 84, 7–17 (1989).

23. Lozoff, B. et al. Iron Deficiency Anemia and Iron Therapy Effects on Infant Developmental Test Performance. Pediatrics 79, 981–995 (1987).

24. Pala, E., Erguven, M., Guven, S., Erdogan, M. & Balta, T. Psychomotor Development in Children with Iron Deficiency and Iron-Deficiency Anemia. Food Nutr. Bull. 31, 431–435 (2010).

25. Algarín, C. et al. Iron-deficiency anemia in infancy and poorer cognitive inhibitory control at age 10 years. Dev. Med. Child Neurol. 55, 453–458 (2013).

26. Lukowski, A. F. et al. Iron deficiency in infancy and neurocognitive functioning at 19 years: evidence of long-term deficits in executive function and recognition memory. Nutr. Neurosci. 13, 54–70 (2010).

27. Roncagliolo, M., Garrido, M., Walter, T., Peirano, P. & Lozoff, B. Evidence of altered central nervous system development in infants with iron deficiency anemia at 6 mo: delayed maturation of auditory brainstem responses. Am. J. Clin. Nutr. 68, 683–690 (1998).

28. Algarín, C., Peirano, P., Garrido, M., Pizarro, F. & Lozoff, B. Iron Deficiency Anemia in Infancy: Long-Lasting Effects on Auditory and Visual System Functioning. Pediatr. Res. 53, 217–223 (2003).

29. Steen, R. G., Xiong, X., Mulhern, R. K., Langston, J. W. & Wang, W. C. Subtle brain abnormalities in children with sickle cell disease: Relationship to blood hematocrit. Ann. Neurol. 45, 279–286 (1999).

30. Radlowski, E. C. & Johnson, R. W. Perinatal iron deficiency and neurocognitive development. Front. Hum. Neurosci. 7, 585 (2013).

31. Wu, Y. et al. The Potential Role of Ferroptosis in Neonatal Brain Injury. Front. Neurosci. 13, 115 (2019).

32. Hare, D. J. & Double, K. L. Iron and dopamine: a toxic couple. Brain J. Neurol. 139, 1026– 1035 (2016).

33. Darki, F., Nemmi, F., Möller, A., Sitnikov, R. & Klingberg, T. Quantitative susceptibility mapping of striatum in children and adults, and its association with working memory performance. NeuroImage 136, 208–214 (2016).

34. Carpenter, K. L. H. et al. Magnetic susceptibility of brain iron is associated with childhood spatial IQ. NeuroImage 132, 167–174 (2016).

35. Hect, J. L., Daugherty, A. M., Hermez, K. M. & Thomason, M. E. Developmental variation in regional brain iron and its relation to cognitive functions in childhood. Dev. Cogn. Neurosci. 34, 18–26 (2018).

36. Zhang, Y. et al. Neonate and infant brain development from birth to 2 years assessed using MRI-based quantitative susceptibility mapping. NeuroImage 185, 349–360 (2019).

37. Ning, N. et al. Spatiotemporal variations of magnetic susceptibility in the deep gray matter nuclei from 1 month to 6 years: A quantitative susceptibility mapping study. J. Magn. Reson. Imaging 49, 1600–1609 (2019).

38. Thomas, E. et al. Newborn amygdala connectivity and early emerging fear. Dev. Cogn. Neurosci. 37, 100604 (2019).

39. Rasmussen, J. M. et al. A novel maturation index based on neonatal diffusion tensor imaging reflects typical perinatal white matter development in humans. Int. J. Dev. Neurosci. 56, 42– 51 (2017).

40. Rasmussen, J. M. et al. Maternal Interleukin-6 concentration during pregnancy is associated with variation in frontolimbic white matter and cognitive development in early life. NeuroImage 185, 825–835 (2019).

41. Rasmussen, J. M. et al. Maternal free fatty acid concentration during pregnancy is associated with newborn hypothalamic microstructure in humans. Obesity 30, 1462–1471 (2022).

42. Hegeman, D. J., Hong, E. S., Hernández, V. M. & Chan, C. S. The external globus pallidus: progress and perspectives. Eur. J. Neurosci. 43, 1239–1265 (2016).

43. Hannah, R. & Aron, A. R. Towards real-world generalizability of a circuit for action-stopping. Nat. Rev. Neurosci. 22, 538–552 (2021).

44. Smith, K. S., Tindell, A. J., Aldridge, J. W. & Berridge, K. C. Ventral pallidum roles in reward and motivation. Behav. Brain Res. 196, 155–167 (2009).

45. Nambu, A., Kaneda, K., Tokuno, H. & Takada, M. Organization of Corticostriatal Motor Inputs in Monkey Putamen. J. Neurophysiol. 88, 1830–1842 (2002).

46. Yu, R., Liu, B., Wang, L., Chen, J. & Liu, X. Enhanced Functional Connectivity between Putamen and Supplementary Motor Area in Parkinson’s Disease Patients. PLOS ONE 8, e59717 (2013).

47. Postuma, R. B. & Dagher, A. Basal Ganglia Functional Connectivity Based on a Meta-Analysis of 126 Positron Emission Tomography and Functional Magnetic Resonance Imaging Publications. Cereb. Cortex 16, 1508–1521 (2006).

48. Groman, S. M. et al. Monoamine Levels Within the Orbitofrontal Cortex and Putamen Interact to Predict Reversal Learning Performance. Biol. Psychiatry 73, 756–762 (2013).

49. Haber, S. N. Corticostriatal circuitry. Dialogues Clin. Neurosci. 18, 7–21 (2016).

50. Tian, M. et al. Morphological Development of the Human Fetal Striatum During the Second Trimester. Cereb. Cortex bhab532 (2022) doi:10.1093/cercor/bhab532.

51. Ragozzino, M. E. Acetylcholine actions in the dorsomedial striatum support the flexible shifting of response patterns. Neurobiol. Learn. Mem. 80, 257–267 (2003).

52. Eagle, D. M. & Robbins, T. W. Inhibitory Control in Rats Performing a Stop-Signal Reaction-Time Task: Effects of Lesions of the Medial Striatum and d-Amphetamine. Behav. Neurosci. 117, 1302–1317 (2003).

53. Collins, P., Wilkinson, L. S., Everitt, B. J., Robbins, T. W. & Roberts, A. C. The effect of dopamine depletion from the caudate nucleus of the common marmoset (Callithrix jacchus) on tests of prefrontal cognitive function. Behav. Neurosci. 114, 3–17 (2000).

54. Grahn, J. A., Parkinson, J. A. & Owen, A. M. The cognitive functions of the caudate nucleus. Prog. Neurobiol. 86, 141–155 (2008).

55. Alexander, G. E., DeLong, M. R. & Strick, P. L. Parallel organization of functionally segregated circuits linking basal ganglia and cortex. Annu. Rev. Neurosci. 9, 357–381 (1986).

56. Lehéricy, S. et al. Diffusion tensor fiber tracking shows distinct corticostriatal circuits in humans. Ann. Neurol. 55, 522–529 (2004).

57. Smyser, C. D., Snyder, A. Z. & Neil, J. J. Functional connectivity MRI in infants: Exploration of the functional organization of the developing brain. NeuroImage 56, 1437– 1452 (2011).

58. Shi, F., Salzwedel, A. P., Lin, W., Gilmore, J. H. & Gao, W. Functional Brain Parcellations of the Infant Brain and the Associated Developmental Trends. Cereb. Cortex 28, 1358–1368 (2018).

59. Molloy, M. F. & Saygin, Z. M. Individual variability in functional organization of the neonatal brain. NeuroImage 253, 119101 (2022).

60. Bethlehem, R. a. I., et al. Brain charts for the human lifespan. Nature 604, 525–533 (2022).

61. Seger, C. A. How do the basal ganglia contribute to categorization? Their roles in generalization, response selection, and learning via feedback. Neurosci. Biobehav. Rev. 32, 265–278 (2008).

62. Deen, B. et al. Organization of high-level visual cortex in human infants. Nat. Commun. 8, 13995 (2017).

63. Cabral, L., Zubiaurre-Elorza, L., Wild, C. J., Linke, A. & Cusack, R. Anatomical correlates of category-selective visual regions have distinctive signatures of connectivity in neonates. Dev. Cogn. Neurosci. 58, 101179 (2022).

64. Holland, D. et al. Structural Growth Trajectories and Rates of Change in the First 3 Months of Infant Brain Development. JAMA Neurol. 71, 1266–1274 (2014).

65. Lyall, A. E. et al. Dynamic Development of Regional Cortical Thickness and Surface Area in Early Childhood. Cereb. Cortex 25, 2204–2212 (2015).

66. Raznahan, A. et al. How does your cortex grow? J. Neurosci. Off. J. Soc. Neurosci. 31, 7174–7177 (2011).

67. Gilmore, J. H., Knickmeyer, R. C. & Gao, W. Imaging structural and functional brain development in early childhood. Nat. Rev. Neurosci. 19, 123–137 (2018).

68. Raznahan, A. et al. Longitudinal four-dimensional mapping of subcortical anatomy in human development. Proc. Natl. Acad. Sci. 111, 1592–1597 (2014).

69. Hammerslag, L. R. & Gulley, J. M. Sex differences in behavior and neural development and their role in adolescent vulnerability to substance use. Behav. Brain Res. 298, 15–26 (2016).

70. Society, C. P. Iron requirements in the first 2 years of life | Canadian Paediatric Society. https://cps.ca/en/documents//position//iron-requirements/.

71. Yuan, X. et al. Iron deficiency in late pregnancy and its associations with birth outcomes in Chinese pregnant women: a retrospective cohort study. Nutr. Metab. 16, 30 (2019).

72. Blüml, S. et al. Metabolic Maturation of the Human Brain From Birth Through Adolescence: Insights From In Vivo Magnetic Resonance Spectroscopy. Cereb. Cortex N. Y. NY 23, 2944– 2955 (2013).

73. Smart, J. L., Dobbing, J., Adlard, B. P., Lynch, A. & Sands, J. Vulnerability of developing brain: relative effects of growth restriction during the fetal and suckling periods on behavior and brain composition of adult rats. J. Nutr. 103, 1327–1338 (1973).

